# TGF-β1 drives Th9 but not Treg cells upon allergen exposure

**DOI:** 10.1101/2021.08.18.456797

**Authors:** Stephanie Musiol, Francesca Alessandrini, Constanze A. Jakwerth, Adam M. Chaker, Evelyn Schneider, Ferdinand Guerth, Ileana Ghiordanescu, Julia T. Ullmann, Josephine Kau, Mirjam Plaschke, Stefan Haak, Thorsten Buch, Carsten B. Schmidt-Weber, Ulrich M. Zissler.

**Author notes:** **Corresponding author:** Prof. Dr. Carsten B. Schmidt-Weber, PhD Center of Allergy & Environment (ZAUM), Technical University of Munich and Helmholtz Center Munich, Biedersteiner Str. 29, 80802 Munich, Germany. equal contributions (Stephanie Musiol, Francesca Alessandrini, Thorsten Buch, Carsten B. Schmidt-Weber, Ulrich M. Zissler.). **Author contribution:** SM, FA, ES FG IG JTU JK MP, SH TB CSW UMZ AMC CAJ CSW, FA, UMZ, TB, SH and AMC developed the study and experimental layout. SM, FA, ES realized experimental disease models and measurements of murine parameters.FA, SH and TB supported the study on ethical permissions and funding. AMC organized study part including human ethical approval and together with JK, MP the management of patient visits, patient information and sampling. Human samples were analyzed by IG, JTU, JK, MP, FG and UMZ. The manuscript was written by CSW, TB, FA and UMZ.

## Abstract

TGF-β1 is known to have a pro-inflammatory impact by inducing Th9 cells, while it also induces anti-inflammatory Treg cells (Tregs). In the context of allergic airway inflammation (AAI) its dual role can be of critical importance in influencing the outcome of the disease. Here we demonstrate that TGF-β acts in AAI by driving effector T cells into Th9 cells, while Tregs differentiate independently. Induction of experimental AAI and airway hyperreactivity in a mouse model with inducible genetic ablation of the TGFβ-receptor 2 (TGFBR2) on CD4^+^T cells significantly reduced the disease phenotype. Further, it blocked the induction of Th9 cell frequencies, but increased Treg cells. To translate these findings into a human clinically relevant context, Th9 and Treg cells were quantified both locally in induced sputum and systemically in blood of allergic rhinitis and asthma patients with or without allergen-specific immunotherapy (AIT). Natural allergen exposure induced local and systemic Th2, Th9 cell and reduced Tregs, while therapeutic allergen exposure by AIT suppressed Th2 and Th9 cell frequencies along with TGF-β and IL-9 secretion. Altogether, these findings support that neutralization of TGF-β represents a viable therapeutic option in allergy and asthma, not posing the risk of immune dysregulation by impacting Tregs.

## 1. INTRODUCTION

The three isoforms of transforming growth factor-β (TGF-β) TGF-β1, TGF-β2 and TGF-β3 in mouse and human, encoded by separate genes, are involved in a plethora of biological processes during development, lineage commitment, wound healing, proliferation, migration and survival of cells ^1^. Each isoform and the TGF-β receptors (TGF-βR) are expressed in specific and temporal patterns making their functions strongly context-dependent for various tissues and cell types ^2^. Within the immune system TGF-β signaling was found to have essential roles in the T cell, B cell and phagocyte compartments, among others. It is essential for the development of regulatory T (Treg) cells, for differentiation and lineage commitment of Interleukin-17 producing T helper (Th17) cells (mouse only), of follicular T helper (Tfh) cells (human only) and potentially of IL-9 producing T helper (Th9) cells ^3, 4^. The various outcomes of TGF-β signaling are achieved through combinatorial sensing of additional cytokines (such as IL-6, IL-1, IL21 in combination with TGF-β for Th17 induction), hence result from specific local cytokine milieus ^5^. Allergic diseases are characterized by an uncontrolled immune reaction towards harmless environmental antigens to which the body is exposed either via airways, as seen in allergic rhinitis (AR) and allergic asthma (AA), via skin (atopic dermatitis), gastrointestinal tract (food allergy) or by systemic exposure (anaphylaxis). In healthy individuals, allergen exposure is tolerated. Loss of T cell tolerance towards environmental antigens is a prerequisite for initiation of an allergic reaction and can lead to activation Th2 cells, humoral (IgE) effector mechanisms as well as infiltration of inflammatory cells at the site of allergen exposure. At these tissue sites, increased mucus production, smooth muscle and airway epithelial cell activation ^6^ are observed, subsumed as airway hyperreactivity (AHR).

In airway inflammation TGF-β is involved in tissue remodeling ^7^, yet all isoforms are expressed throughout the normal lung, including expression by bronchial epithelium, macrophages, vascular endothelium, smooth muscle and fibroblasts ^8, 9^. TGF-β is also a core inducer of the epithelial-mesenchymal transition process during fibrotic remodeling of airways ^10^. Both, TGF-β1 and TGF-β2 have been shown to be increasingly expressed during airway hyperreactivity (AHR), especially in eosinophils ^11^ and macrophages ^12^; in case of TGF-β2, in epithelium ^13^ and neutrophils. The amount of TGF-β1 in bronchoalveolar lavage (BAL) is increased in AA and both TGF-β1 and −2 levels are increased upon segmental allergen challenge of the lung ^14^. Similar observations have been made in the allergic airway inflammation (AAI) model(s). TGF-β is not only part of the regulatory mechanisms of Tregs; they are themselves developmentally dependent on sensing TGF-β ^15^. Tregs can keep the potentially pathogenic IL-4-producing Th2 cells under control and thus avoid AAI ^16^. The production of IL-4 by Th2 cells is itself not pathogenic, as it is also observed in healthy individuals ^17^. In contrast, IL-9-producing T helper cells (Th9) were recently identified as critical subset in AA ^18^. In fact, increased IL-9-secretion was observed in BALF and lung tissue of AA patients ^19^. IL-9 was described to induce mucus production by epithelial cells ^20^, support *de novo* mast cell generation and their proliferation *in situ* ^20^. It further serves as chemoattractant for recruitment of inflammatory cells to sites of inflammation ^21^.

The origin of Th9 cells in AA is still unclear. However, TGF-β-dependent ^22^ and –independent ^23, 24^ mechanisms of induction were described. Recent studies showed that also environmental sensor mechanisms via the mTOR pathway and Foxo1/3 can trigger Th9 cell differentiation ^25^. TGF-β primes together with IL-4 the production of IL-9 in PPARγ^+^ Th2 cells ^26^, as also shown for the induction of Th17 and Treg cells ^27^. In contrast to Th17 cells, the Vitamin A metabolite retinoic acid suppresses Th9 differentiation ^28^.

Different mouse models have laid the basis of our understanding of the role of TGF-β *in vivo*. A complete absence of TGF-β signaling from the T cell compartment was found to result in profound and detrimental autoimmune pathologies ^29^. Yet, the exact delineation of TGF-β in these pathologies remain an ongoing task, also due to the pleiotropic function of TGF-β signaling in different cell types. Among those, Th9 cells are in focus since TGF-β is known to be critical for their development and Th9 counts directly correlate with AHR severity ^30^, while IL-9 neutralization relieves local inflammation ^31^.

Therefore, we wanted to test whether presence of TGF-β signaling was indeed a necessary requirement for the *in vivo* development of Th9 cells in the context of AAI. We used our system for inducible ablation of TGF-βRII in Th cells for selectively interrogating the role of TGF-β signaling in a mouse model of AAI. Confirming a selective role of TGF-β signaling for development of Th9 cells in AAI, and rejecting its involvement in Treg homeostasis in our model we set out to confirm this pathway in human rhinitis patients. We took advantage of induced sputum as a non-invasive window to lung pathology in allergic patients and in context of allergen-specific immunotherapy (AIT). We confirmed opposite regulation of TGF-β with respect to IL-4 and IL-9 by disease and treatment, however we found no influence of local TGF-β signaling on Treg cells. These findings are of particular importance as Th9 cells are uniquely important for lower airway inflammation compared to other type-2 diseases.

## RESULTS

### Reduced airway hyperreactivity and lung cell infiltration in allergic iCD4TGFBR2 mice

To address the question whether TGF-β is driving Th9 or Treg cells upon allergen exposure we used OVA-induced AAI in a mouse model. We compared presence versus absence of TGF-β signaling by use of tamoxifen-facilitated Cre-loxP-mediated deletion of TGFBR2 in CD4^+^ cells after the sensitization phase. These iCD4TGFBR2 ^32^ were compared to wildtype (WT) C57BL/6/J animals (Fig.1). Twenty-four hours after the last allergen challenge we observed decreased AHR to methacholine in absence of TGF-β signaling (Fig. 1B). TGFBR2 ablation significantly decreased the number of leukocytes, specifically eosinophils, neutrophils and lymphocytes but not macrophages in BAL, with the most prominent change in the fraction of eosinophils (Fig. 1C and D). PAS-staining of lungs revealed decreased perivascular and peribronchiolar inflammatory cell infiltration (Fig. 1 E-H) and mucus hypersecretion following AAI upon TGFBR2 ablation, (Fig. 1 E, I), confirming the ameliorated clinical outcome. These local changes to the severity of disease in our AAI model by abrogation of TGF-β signaling in CD4^+^ T cells were reflected in serum, in which the levels of total IgE (Fig.1 F) and OVA-specific IgE (Fig.1 G) were drastically reduced. Without induction of AAI, induced Th cell specific TGFBR2-ablation did not result in any of the above-described outcomes, hence highlighting the critical role of TGF-β signaling during the challenge phase of established AAI.

**Figure 1.**
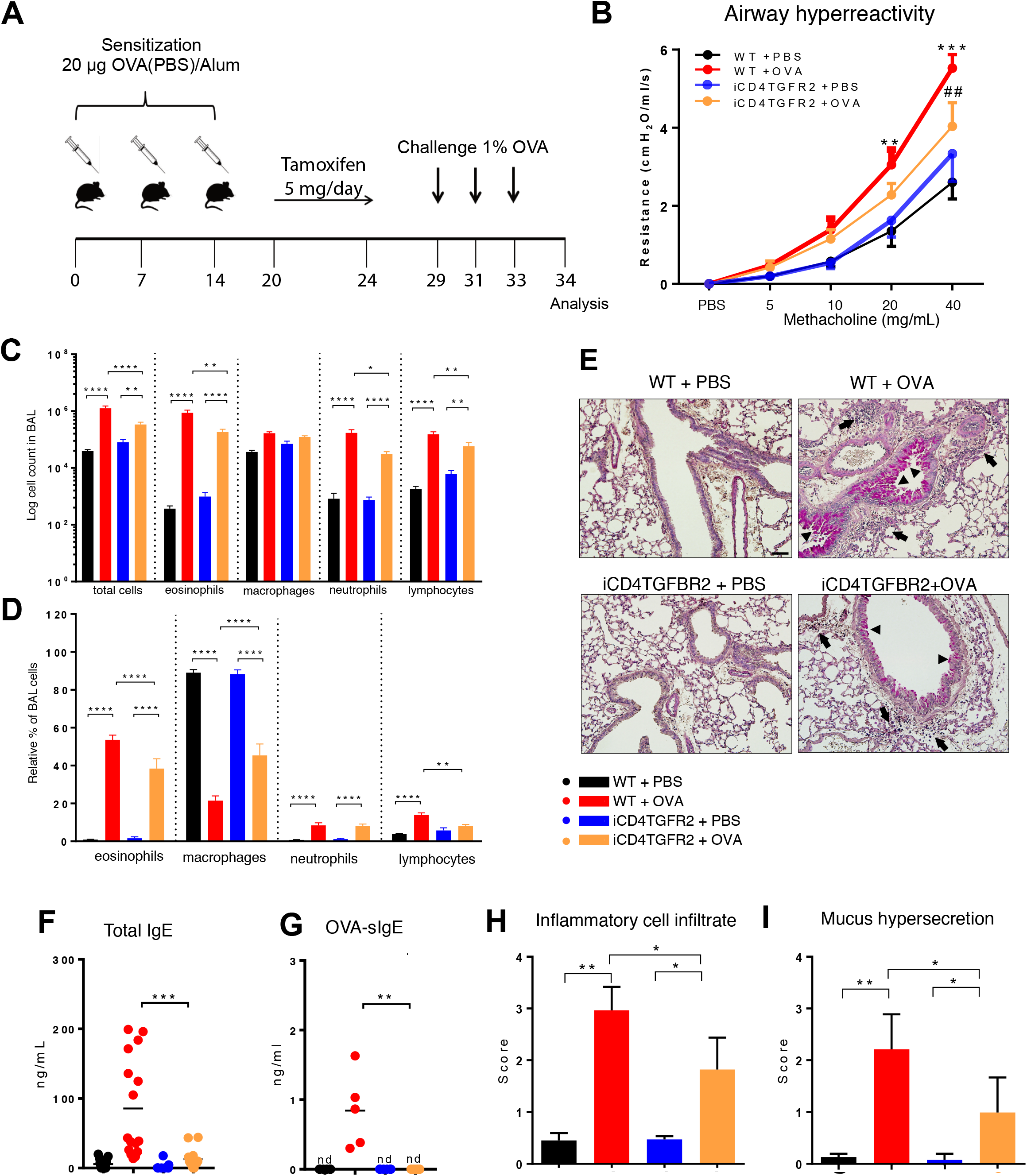
Impact of TGFBR2 ablation on lung inflammatory infiltrate and AHR. **(A)** Scheme of AAI induction, challenge and tamoxifen treatment. **(B)** Airway hyperreactivity measured in intubated, mechanically ventilated animals following methacholine provocation. n= 5 - 9/group; **≤0.01; ***≤0.01 vs WT+PBS; ^##^p≤0.01 vs WT+OVA at same methacholine concentrations (two-way analysis of variance (ANOVA) with Bonferroni’s post-hoc test). **(C)** BAL total cell number and **(D)** relative population size of mice from the four experimental groups. **(E)** PAS-staining of lung sections from the four experimental groups. Arrows: inflammatory infiltrate; arrowheads: mucus hypersecretion; scale bar: 50 μm. **(F-G)** Levels of total and OVA-specific Immunoglobulin E (tIgE: n=11-16/group; OVA-sIgE: n=5/group) measured in plasma samples (two-tailed Mann-Whitney U test). **(H-I)** Histological scores of **(H)** Inflammatory cell infiltrates and **(I)** mucus production (mean ± SD; n=5/group). C–I: representative of three independent experiments each with 5 mice per group.

### Cytokine response upon ablation of TGFBR2 in CD4^+^T cells during AAI

To better understand how abrogation of TGF-β signaling before the challenge phase was ameliorating AAI, we determined cytokine levels in BAL fluid. As expected, AAI resulted in local increase of all measured cytokines in WT animals (Fig. 2A, Table S1). In absence of TGFBR2 signals within Th cells we observed significantly less IL-4, −5, −6, −9, −17A, IFN-γ (Fig. 2A, Tab. S1A) and TNF-α (Fig.S1A) while CXCL-1, −2 and CCL-2 and −3 showed the same trend, but did not reach statistical significance (Fig.S1A). Similarly, serum levels of IL-9 appear lower following TGFBR2 ablation in allergic animals (Fig. S1C), however did not reach significance. On the contrary, BAL IL-2, IL-10 and IL-13 levels remained unaffected (Fig.2A).

**Figure 2.**
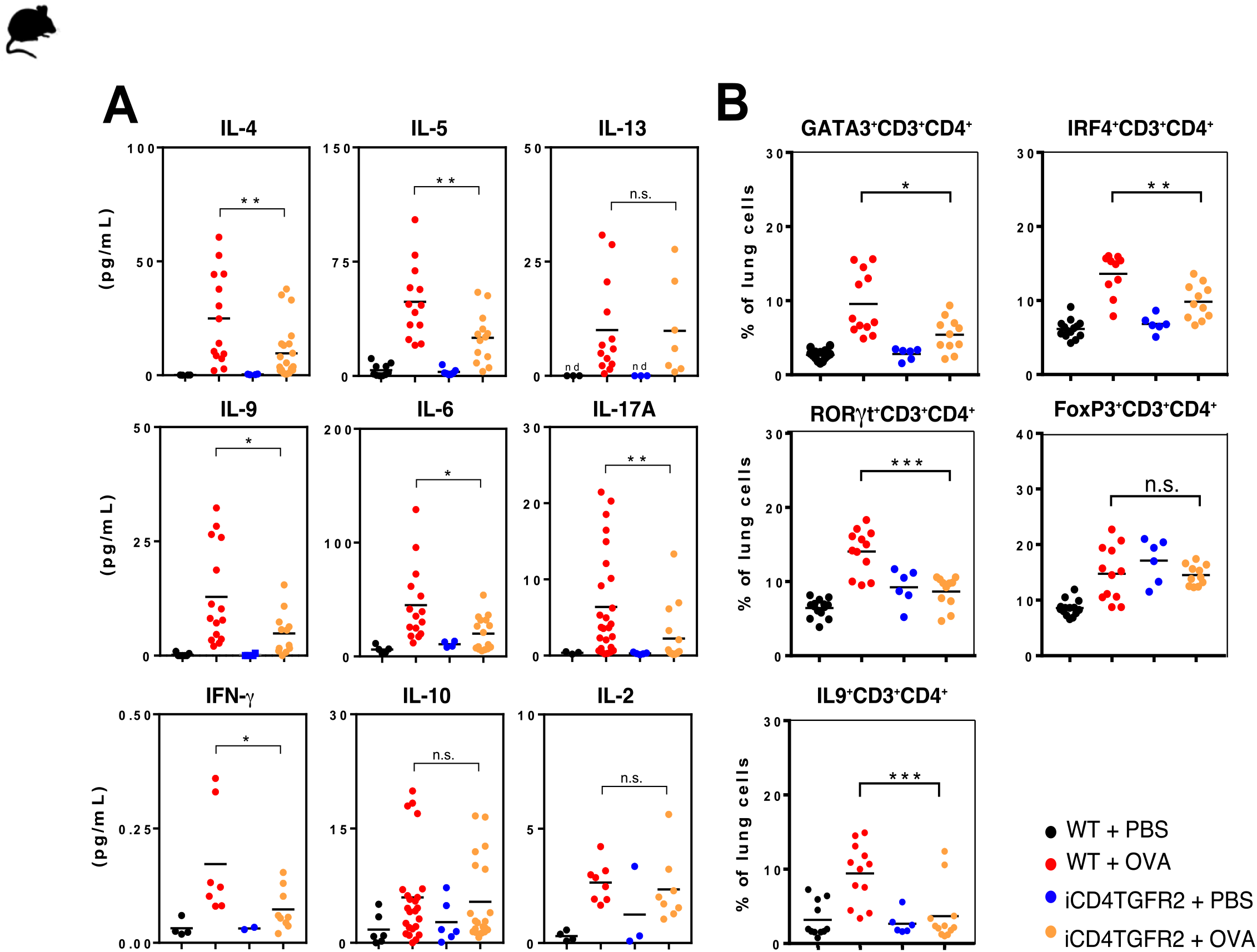
Impact of TGFBR2 ablation on lung cytokine production and T cell subsets. **(A)** BAL fluid of the indicated experimental groups was analyzed using electrochemiluminescence assay. Each data point represents an individual mouse. Data is compiled from one or two independent experiments (n=5-15/group). *p<0.05, **p<0.01, ***p<0.001, ****p<0.0001 (t-test). **(B)** Flow cytometric analysis of lungs focusing on distinct T cell subsets, performed 24h after the last OVA-challenge. Percentages of Th2 (GATA3^+^), Th9 (IRF4^+^; IL-9^+^), Th17 (RORγt^+^) and Treg (FoxP3^+^) cells within the CD3^+^CD4^+^T cell population in non-allergic WT+PBS and iCD4TGFBR2+PBS mice and allergic WT+OVA and iCD4TGFBR2+OVA mice were assessed. Each data point represents an individual mouse. Data are representative of two independent experiments (n=6-13/group). *p<0.05, **p<0.01, ***p<0.001, ****p<0.0001 (two-tailed Mann-Whitney U test).

Analysis of the specific cytokine response by splenocytes after *in vitro* restimulation with OVA revealed reduced production of IL-2 and IL-5 after TGFBR2 ablation, while IL-4 and IL-6 were slightly decreased (Fig. S1D-E). The specificity of immunological effects is supported by the unchanged TNF-α secretion (Fig. S1E).

We next confirmed that the source of the BAL fluid and serum cytokines described above were lung T cells. We found decreased frequencies of GATA3^+^ Th2 cells, IRF4^+^ or IL-9^+^ Th9 cells and RORγt^+^Th17 following TGFBR2 ablation in allergic animals (Fig.2B). Independent of AAI induction, the TGFBR2 ablation led to increased fraction of FOXP3^+^ Tregs in the lung (Fig.2). Unexpectedly, we also found the fraction of lung CD8^+^ T cells to be increased in absence of TGF-β signaling in CD4^+^ T cells, while CD4 T cells themselves were unchanged (Fig. S1F). γδ T-cell percentages remained unchanged (Fig.S1F). Taken together, abrogated TGF-β signaling resulted in reduced responses of the Th2, Th9, and Th17 lineages during AAI while Treg cells expanded independently of AAI.

### Gene network and transcriptional analysis

To further understand T cell differentiation in allergy, a gene interaction network was assembled (Fig. 3A). Key transcription factors for Th9 cells are IRF-4 and PU.1, whereby IRF-4 is known to synergize with PU.1 ^33^. In fact, *Pu.1* appears to be essential for *IL9* expression ^34^, and IRF4 can interact with SMAD-3 to promote *Il9* expression ^35^, while the IL-4 and TGF-β inducible factors BLIMP-1 (not shown) inhibits IL-9. IRF4 also facilitates expression of the IL-17 gene ^36^. Th9 and Th17 are not only linked by IRF4 and SMAD3, but also FOXO1 and FOXO2 are regulating both T cell subsets ^37^. To assess the different components of this network we performed expression analysis on CD4^+^ T cells from lungs of animals of our four experimental groups. Allergic inflammation enhanced expression level of *Il9, Nfat5, Foxo1* and *Foxo3*, while PU.1, *Sgk1* and *BATF* remained unchanged (Fig. 3B). Ablation of TGFBR2 decreased expression of *Il9*, *Pu.1*, *Foxo1*, *Foxo3* and *Nfat5* in diseased animals. The current knowledge of gene regulation networks of Th9 cells is overlapping or linked with Th2 and Th17 genes and some of these factors are connected to TGF-β signaling (Fig. 3C). To obtain a schematic overview on gene regulation we assessed mRNA levels of these factors in the lungs with and without induction of AAI. Allergic inflammation enhanced expression level of some Th17, Treg and Th9 factors including IRF4 (Fig.3C; red), while the Th9 transcription factor PU.1 remained unaffected (left). TGFBR2 ablation decreased the expression of Th17 and Th9 factors, while Th2 and Treg factors staid unchanged (Fig. 3C) compared to WT (left). Interestingly, the IRF4 increase induced by allergic inflammation (left) is not affected by TGFBR2 ablation (right) despite the relationships of IRF4 to the TGF-β signaling.

**Figure 3.**
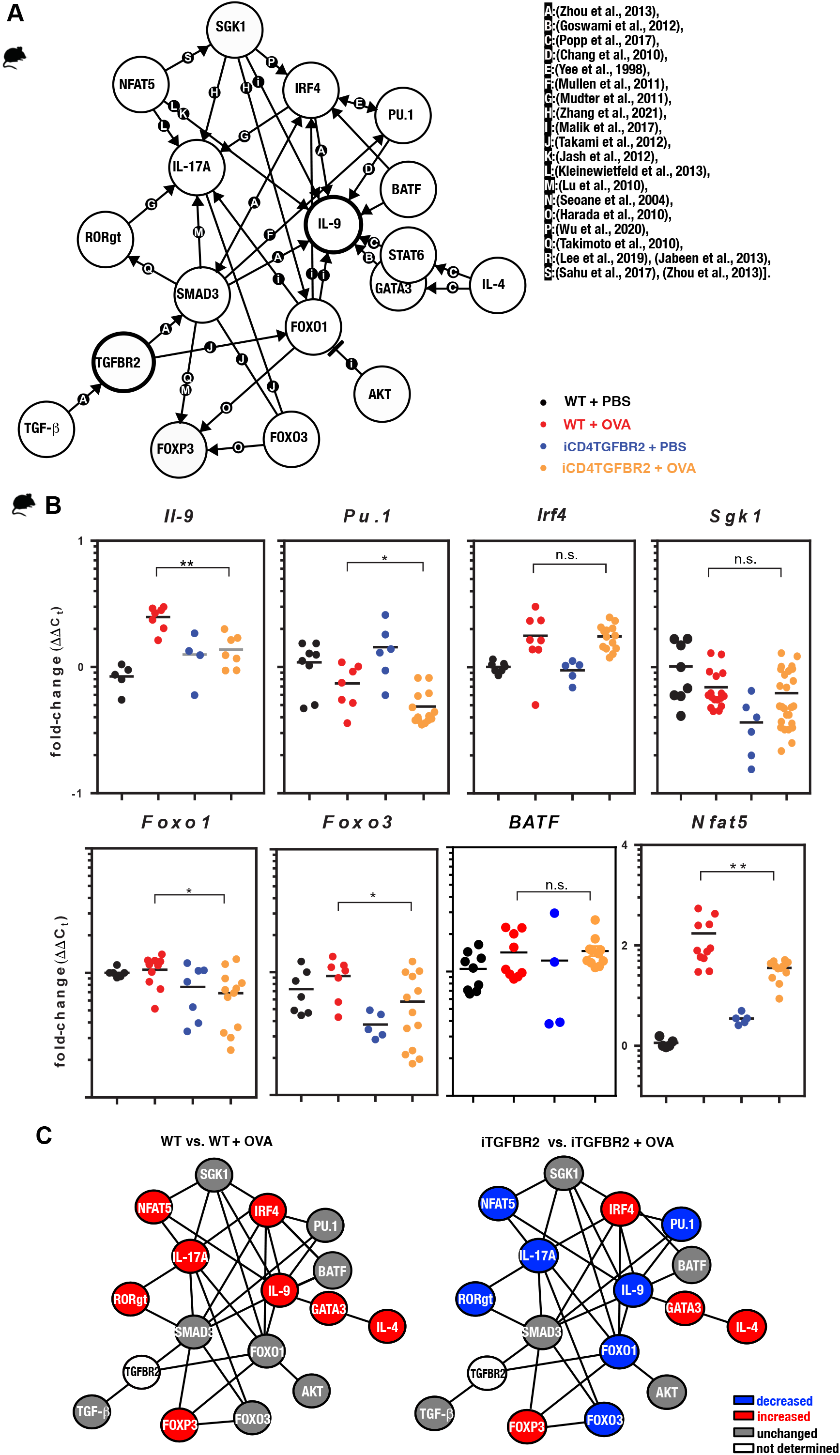
**(A)** Illustration of the complex interaction of genes in the regulation of T cell differentiation in allergy. The current knowledge was compiled in a gene regulation network of TH9 cells, which overlaps with T_H_2 and T_H_17 networks. The references for the network are as follows: [a:(Zhou et al., 2013), b:(Goswami et al., 2012), c:(Popp et al., 2017), d:(Chang et al., 2010), e:(Yee et al., 1998), f:(Mullen et al., 2011), g:(Mudter et al., 2011), h ^50^, i:(Malik et al., 2017), j:(Takami et al., 2012), k:(Jash et al., 2012), l:(Kleinewietfeld et al., 2013), m:(Lu et al., 2010), n:(Seoane et al., 2004), o:(Harada et al., 2010), p:(Wu et al., 2020), q:(Takimoto et al., 2010), r:(Lee et al., 2019), (Jabeen et al., 2013), s:(Sahu et al., 2017), (Zhou et al., 2013)]. **(B)** CD4^+^ T cells were isolated from digested lungs of non-allergic and allergic WT and iCD4TGFBR2 mice and analyzed for IL-9 expression as well as T_H_9 and T_H_2 relevant transcription factors and signal molecules using quantitative RT-PCR. Expression levels of *Il9, Pu.1, Irf4, Sgk1, Foxo1, Foxo3*, *Batf* and *Nfat5* were normalized to *Gapdh* house-keeping gene and relative changes were represented as 2^−ΔΔCT^ (ΔΔC_T_=ΔC_T_−ΔC_Control_). Data is compiled from two independent experiments (n= 5-13/group). *p<0.05, **p<0.01, ***p<0.001 (t-test). **(C)** Comparisons of gene regulation network of T cells. Blue indicates reduced gene expression following OVA-induced AAI, while red indicates increased and gray unchanged gene expression of the respective factor.

Taken together, ablation of TGFBR2 seems to affect the transcriptional network of Th17 and Th9 cells, leaving the Th2 and Treg networks largely untouched.

### Effect of AIT on TGF-β and T cell lineage specific cytokines in induced sputum of allergic patients

In our AAI mouse model, we observed very specific effects of absence of TGF-β signaling on cytokine production and the presence of respective T cell lineages in the lung. We proved the influence of TGF-β on local allergic airway reactions with a focus on established AAI. To confirm the relevance of these results for human allergy, we next investigated immune cells of the lower airways of rhinitis patient without and with asthma comorbidity in the context of AIT. Our cross-sectional cohort consisted of 26 healthy controls and 38 allergic rhinitis patients (AR), 19 of these with asthma comorbidity (AA; table S4). Ten AR patients and nine AA patients received AIT, while nine untreated AR and ten untreated AA were assigned to the untreated groups. In induced sputum samples from our patient group we observed that AIT reduced back to baseline the elevated levels of TGF-β found both for AR and AA patients *in season* of natural pollen exposure (May-July, Fig. 4A, table S5). Hence, allergic patients under AIT reveal a situation similar to the one artificially obtained by TGFBR2 ablation in the mouse model. A reduction back to baseline was also observed for IL-4, IL-5, and IL-13 as well as IL-9, thus the classical cytokines of Th2 and Th9 cells respectively (Fig. 4B, table S5). The levels of IL-2, IL-6, IL-10 and IFN-γ, hence pan-T cell, inflammatory, Treg and Th1 cytokines, respectively, showed the opposite reaction with higher sputum levels upon AIT (Fig. 4B). While the effect sizes were smaller, these observations were confirmed *out of seaso*n (October-January, Fig. S2A). We also observed that sputum levels of TGF-β1 showed a positive correlation with total IgE (r=0.3488, *p*=0.0189; data not shown).

**Figure 4.**
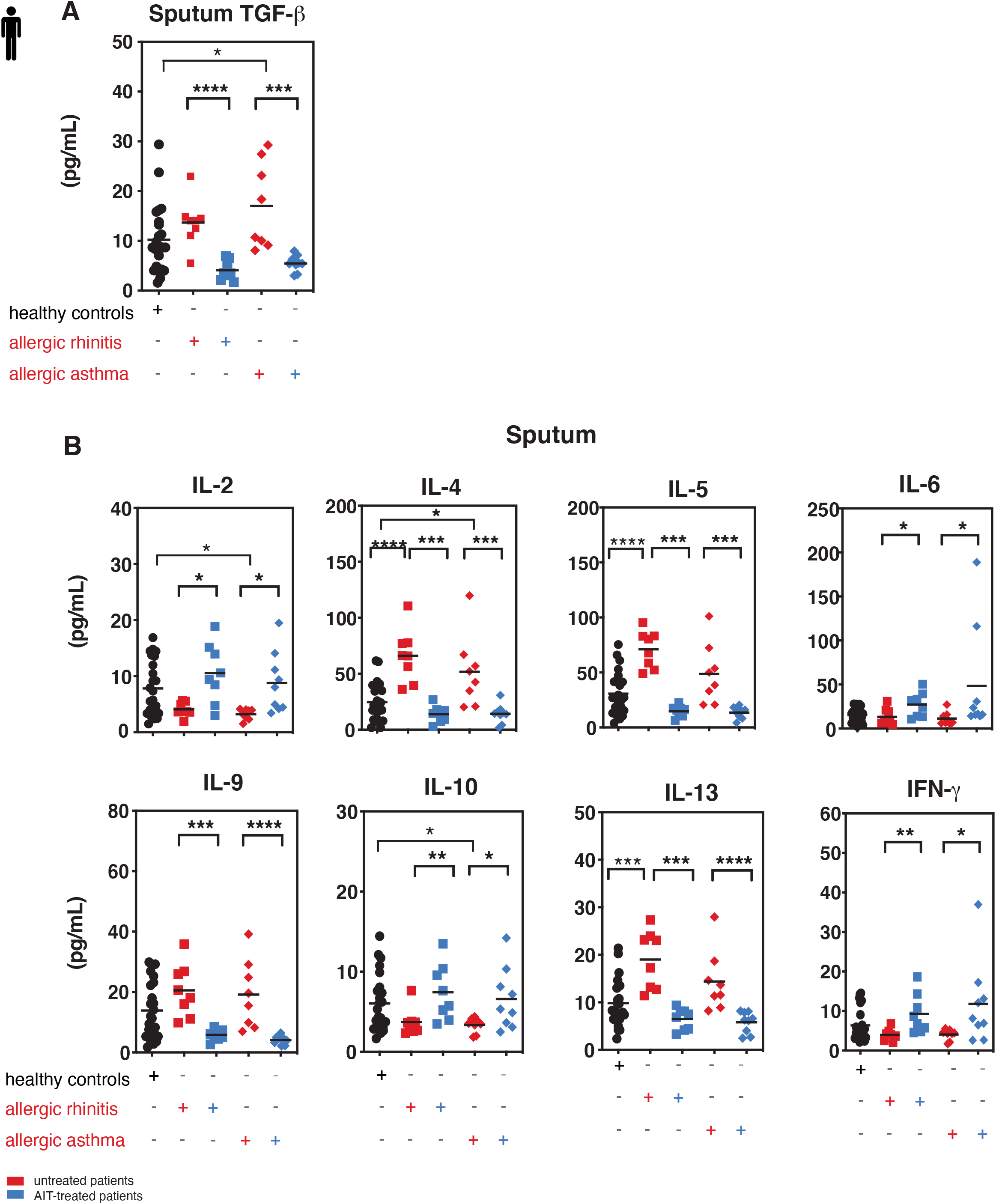
Local TGF-β is decreased along with pro-inflammatory cytokines upon AIT. Levels of selected cytokines in sputum of treated and untreated AR and AA patients and healthy controls. **(A)** Secreted TGF-β cytokine levels in induced sputum of allergic rhinitis and allergic asthma patients receiving (blue colored) or not (red colored) AIT as well as of control individuals were analyzed by LegendPlex. Data presented by individual values and mean. **(B)** Levels of secreted cytokines IL-2, IL-4, IL-5, IL-6, IL-9, IL-10, IL-13 and IFN-γ detected in sputum supernatants of the groups as in (A) assessed by Legendplex. Data presented by individual values and mean. *p<0.05, **p<0.01, ***p<0.001, ****p<0.0001; statistical significance was determined by Kruskal-Wallis tests and only when medians across patient groups varied significantly, multiple single comparisons were performed using two-tailed Mann-Whitney U tests.

Taken together, in AR and AA patient airways TGF-β levels are reduced and Th2 and Th9 cytokines are ameliorated following AIT.

### Decreased Th2 and Th9 and increased Treg cells upon AIT

Since TGF-β has distinct effects dependent on specific tissue context, we quantified the frequency of Th9 cells (TGFBR2^+^IL9^+^CD3^+^CD4^+^), Th2 cells (GATA3^+^ CD3^+^CD4^+^), and Treg cells (FoxP3^+^ CD3^+^CD4^+^) in induced sputum samples and PBMCs by flow cytometry (Fig. 5). *In season*, frequencies of sputum Th2 and Th9 cells were strongly reduced in treated patients compared to untreated patients, both in sputum (Fig. 5C) and in blood (Fig. 5E). Percentages of Treg cells were, however, increased after AIT (Fig. 5C, E). Hence, for all three T cell lineages frequencies were basically normalized by AIT. Similar effects, albeit less pronounced, could be observed *out of season* (October-January) of natural pollen exposure (Fig. 5D, F). Surprisingly, *out of season* the amelioration of Th2 frequencies by AIT in sputum was very small in contrast to the Th9 frequency change (Fig. 5D), while in blood the effect size remained large (Fig. 5F). The size of the Th2 and Th9 subsets in sputum and blood presented only weak or no correlation to serum IgE levels (Fig. 5G, I), whereas the symptom score RQLQ, most relevant as therapy outcome parameter, was strongly positively correlated to the size of both Th subsets (Fig. 5H, J). These results demonstrate that AIT modulates the local and systemic abundance of pro-allergic Th2 and Th9 cell subtypes, while Treg cells and local TGF-β were increased.

**Figure 5.**
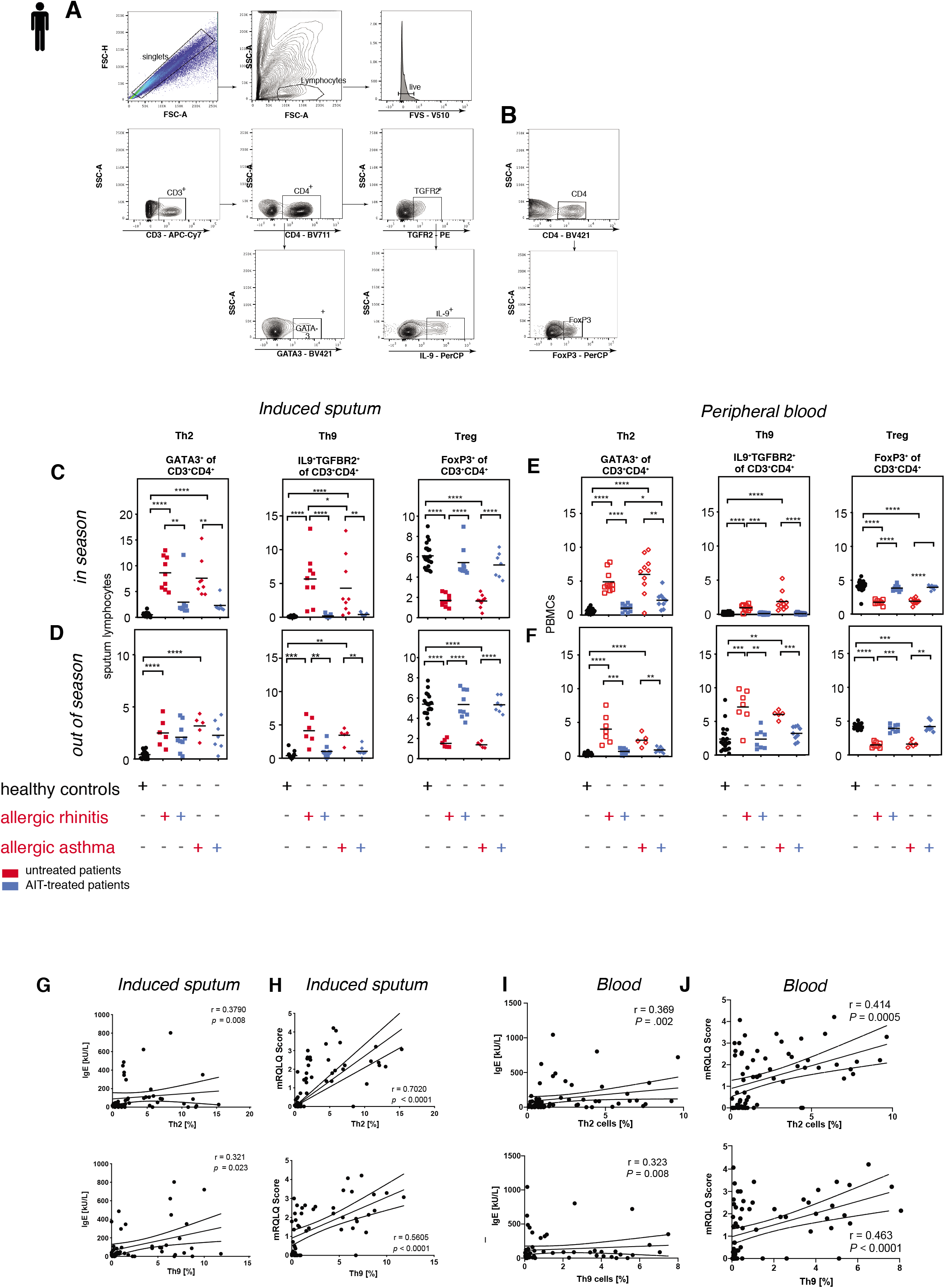
Tregs increase and Th9 as well as Th2 cells decrease upon AIT. T_H_2, T_H_9, and Treg frequencies in sputum of treated and untreated AR and AA patients and healthy controls. **(A and B)** Flow cytometry analysis and gating strategy for T_H_2 and T_H_9 **(A)** and Treg cells **(B)** in induced sputum. **(C-D)** Representative example plots of flow cytometric analysis of sputum derived T cell subpopulation Th2, Th9 and Tregs in healthy subjects, AR and AA patients as described in *in season* **(C)** and *out of season* **(D)**. **(E-F)** Representative example plots of flow cytometric analysis of peripheral blood derived T cell subpopulation Th2, Th9 and Tregs in healthy subjects, AR and AA patients as described in *in season* **(E)** and *out of season* **(F)**. Data presented by single patient values and mean. Statistical significance was determined by Kruskal-Wallis tests, which were performed initially and only when medians across patient groups varied significantly, multiple single comparisons were performed using two-tailed Mann-Whitney U tests. *p<0.05, **p<0.01, ***p<0.001, ****p<0.0001. **(G-H)** Correlation of sputum T_H_2 and T_H_9 cell frequencies with total serum IgE **(G)** and symptom score mRQLQ **(H)**. **(I-J)** Correlation of peripheral blood derived T_H_2 and T_H_9 cell frequencies with total serum IgE **(I)** and symptom score mRQLQ **(J)**. Two-sided Spearman test was used to calculate the correlations.

## 4. DISCUSSION

The TGF-β pathway was so far thought to play contradictory roles in the course of allergic airway disease precluding further consideration as a drug target. Based on this information, addressing the TGF-β pathway therapeutically was thought to lead to reduction of Th9 activity and stalled fibrosis, yet at the expense of a deregulation of the T cell response and beyond. Therefore, our study was aimed at dissecting these contradictory pro- and anti-inflammatory roles of TGF-β with a focus on Th9 and Treg cells at the site of allergic airway inflammation by combining a murine disease model with human patient data.

Since IL-9 and TGF-β 1 are reported to play a similar role in human ^17^ and murine ^38^ allergic inflammation, we started our investigation by using a murine model of AAI with induced TGFBR2-ablation in CD4^+^T cells ^32^ to understand the balance between pro- and anti-inflammatory roles of the TGF-β pathway in detail. We demonstrate that mice with TGFBR2-ablation presented reduced clinical features of AAI, confirmed by strongly ameliorated histological and functional disease parameters. Hereby, the study shows that during AAI TGF-β-signaling in CD4^+^T cells is an important factor for Th9 cell differentiation *in vivo,* while the Treg compartment remained largely unaffected. Th2, Th9 and Th17 cell recruitment upon allergen challenge ^39^ was reduced in absence of TGFBR2 along with respective transcription factors PU.1 (Th9) and GATA3 (Th2). Moreover, the transcription factor FOXO1, which was recently described to be essential for IL-9 production and Th9 cell differentiation, thus promoting airway allergy ^25, 40, 41^, was clearly reduced upon TGFBR2-ablation. This specific impact of TGFBR2-depletion on T cell subsets during AAI was not limited to the respiratory system, but also extended to a systemic response measured in splenocytes. In line with these reduced sizes of pathologically relevant T cell lineages, disease activity was also ameliorated in terms of infiltrating myeloid cells and production of total and specific IgE. In the context of AAI, ablation of TGF-β signaling on T cells had astonishingly no effect on the size of the local and systemic Treg compartment.

Taken together, absence of TGF-β signaling by CD4^+^ T cells after sensitization did not interrupt the immune control by the Treg compartment, while we clearly confirmed a major role for the Th9 and Th2-driven allergic response upon challenge.

Also, in human patients suffering from AR, treatment with AIT shows decreased levels of sputum TGF-β along with the pro-inflammatory type-2 cytokines. We took advantage of this treatment-induced TGF-β-modulation and monitored Treg and Th9 cells. In the AIT treated patients we demonstrate that the seasonally induced levels of Th9/Th2 cytokines were strongly decreased, locally and systemically, while Th1, general inflammatory cytokines as well as the suppressive cytokine IL-10 were increased. These observations were mirrored by the respective population sizes, again locally and systemically. While local sputum cytokine levels were not yet investigated in context of AIT, it was previously shown that AIT controls IL-9 expression in the upper airways ^42^, as well as Th2 frequency in biopsies ^42^ and peripheral blood ^17^. Interestingly, Th2 cells were also detectable in sputum of health individuals both *in-* and *off-season*. This finding confirms our previous studies, in which we reported IL-4 mRNA and secreted IL-4 protein at low, but detectable levels in the airways of healthy individuals ^17^.

While the presence of Tregs was previously detected in sputum ^43^ we were surprised that Tregs constituted the most abundant T cell subpopulation in the sputum of healthy individuals *in* and *off season*. Furthermore, untreated AR and AA patients consistently displayed strongly reduced Tregs frequency compared to healthy individuals even *off-season*. In contrast, the AIT treatment restored Treg cells percentages, locally and systemically, even though TGF-β levels were reduced by AIT. Although the study was not powered to quantify treatment success, the symptom load was reduced by AIT treatment and correlated strongly with the local, and somewhat less with the reduced systemic percentages of Th2 and Th9 cells.

A reduction of Tr1 cells in the context of allergic airway disease has been reported previously and may be represented in our dataset by the reduction in IL-10 in induced sputum of allergic patients ^44^.

Taken together, our data demonstrate that TGF-β acts in allergic airway inflammation by reprogramming T cells into Th9 cells. Since Tregs remain unaffected (mouse) or are even increased (patients with AIT) by reduced levels of TGF-β signaling, targeting of the TGF-β pathway might therefore not only suppress tissue-remodeling processes but also target pro-allergic Th9 cells, without the risk of affecting immune tolerance adversely.

## METHODS

### Animals and murine model of AAI

C57Bl/6J mice, originally obtained from Charles River, and CD4Cre^ERT2^Tgfbr2^fl/fl^ mice (iCD4TGFBR2) were bred at the animal facility at the Institute of Comparative Medicine of the Helmholtz Centre Munich. Generation and characterization of iCD4TGFBR2 mice was described elsewhere ^32^. All mice were co-housed under specific pathogen-free conditions in individually ventilated cages (VentiRack) and fed by standard pellet diet (Altromin Spezialfutter GmbH & Co. KG) and water *ad libitum*. Both female and male mice aged 6-8 weeks were used for the experiments. To induce AAI an established ovalbumin-sensitization model was used ^45^. Briefly, iCD4TGFBR2 and WT C57Bl/6J mice were sensitized by i.p.-injections of 20 μg ovalbumin (OVA; grade V; Sigma-Aldrich, USA) in phosphate buffered saline (PBS) adsorbed to aluminum hydroxide (2.5 mg, ImjectAlum) on days 0, 7 and 14. Non-allergic control mice received the same volume of PBS in alum. The assignment to the two different groups occurred randomly. On days 29, 31 and 33 all mice were aerosol-challenged for 20 minutes with 1% OVA in PBS delivered by a Pari-Boy nebulizer (Pari), ^45^. The experimental protocol is depicted in Fig.1A. Blood samples were taken before sensitization and at the end of the experiment. Animals were sacrificed twenty-four hours after the last OVA challenge. The study was conducted under federal law and guidelines for the use and care of laboratory animals and was approved by the Government of the District of Upper Bavaria and the Animal Care and Use Committee of the Helmholtz Center Munich (approval number: 55.2-1-54-2532-75-2012).

### Tamoxifen treatment

To ablate the subunit 2 of the heterodimeric TGF-β receptor, mice were treated with tamoxifen (TM, tamoxifen-free base, Sigma-Aldrich) after the sensitization phase. Tamoxifen was suspended in 100% ethanol to 1 g/ml, vortexed, and mixed with corn oil (Sigma-Aldrich) to a final concentration of 100 mg/ml. Before *in vivo* administration, the solution was heated to 37°C until it was properly dissolved. On day 20-24, 50 μl (5 mg) tamoxifen per day were administered by intra-gastric gavage in both CD4Cre^ERT2^Tgfbr2^fl/fl^ and WT control mice ^32^.

### Measurement of airway hyperreactivity

AHR to methacholine (Mch; Sigma-Aldrich) was measured 24 h after the last OVA challenge in intubated, mechanically ventilated animals (n = 6–10/group; Data Sciences International (DSI), as previously described ^45^. Briefly, animals were anesthetized by an intraperitoneal injection of Ketamine (100 mg/kg) and Xylazine (5 mg/kg) in PBS. After cannulation of the trachea and starting mechanical ventilation, the animals were challenged with increasing methacholine (Mch) concentrations, using an in-line nebulizer (5 μl Mch solution in PBS delivered for 30 seconds at the following concentrations: 0, 5, 10, 20 and 40 mg/ml). Data were recorded using the FinePoint software v2.4.6 (DSI). The highest values of respiratory system resistance (R) were recorded every 5 seconds during the data recording interval set at 3 min after each Mch level. The heart rate of each animal was continuously monitored using an ECG device connected with three subcutaneous electrodes throughout the entire experiment (DSI).

### Analysis of bronchoalveolar lavage, lung histology and serology

BAL and evaluation of inflammatory cell infiltration were performed as described previously ^45^. Aliquots of cell-free BAL fluid were used to measure cytokines and chemokines via mesoscale technique using two different kits (V-Plex proinflammatory panel 1 mouse kit and V-Plex cytokine panel 1 mouse kit; MesoScaleDiscovery) according to manufacturer’s instructions. Total and ovalbumin-specific IgE were measured in plasma samples taken before and at the end of the experiment by ELISA as described previously ^46^. For lung histology, after BAL, the lungs were excised and the left lobe fixed in 4% buffered formalin and embedded in paraffin. Sections of 4μm thickness were stained with hematoxylin-eosin (H&E) and periodic acid Schiff (PAS). Mucus hypersecretion and inflammatory cell infiltration were graded in a blinded fashion on a scale from 0 to 4 (0=none, 1=mild, 2=moderate, 3=marked, 4=severe), reflecting the degree of the pathological alteration.

### Isolation and analysis of leukocytes from lung tissue

Lungs were excised, cut into small sized pieces and digested in RPMI medium supplemented with 100 μg/ml DNAse (Sigma-Aldrich) and 1 mg/ml Collagenase Type 1A (Sigma-Aldrich) at 37°C. Digested lungs were filtered through a 70 μm cell strainer, pelleted (400 G, 4°C, 5 min) and resuspended in 6 ml 40% percoll in RPMI (v/v) solution, which was underlayed with 4 ml 80% percoll solution (GE Healthcare-Life Sciences, Chicago, IL, USA). Tubes were centrifuged (1600 G, RT, 15 min) with brake set to 0. Lymphocytes were collected from the interphase and analyzed by flow cytometry.

### Isolation and restimulation of splenocytes

Spleens were excised and single cell suspensions were obtained and re-stimulated as previously described ^47^. Cells were washed and re-suspended in complete medium [RPMI 1640 supplemented with 10% FCS, 1% glutamine, 1% penicillin-streptomycin, 1% Na-pyruvate, 1% non-essential amino acids (Gibco, Life Technologies GmbH) and 50μM 2-β-mercaptoethanol (Sigma-Aldrich)], plated in 96-well at a concentration of 2×10^5^ cells/well and cultured for 72 h with medium alone or with OVA V (5 μg/ml; Sigma-Aldrich). Their supernatants were analyzed for cytokine expression using the V-Plex proinflammatory panel 1 mouse kit (MesoScaleDiscovery) according to manufacturer’s instructions.

### Patients

Specimen of 26 healthy controls and 38 AR patients, of whom 19 received AIT (table S4). All subjects completed a Rhinoconjunctivitis Quality of Life mini Questionnaire (mRQLQ), a lung function test and induced sputum induction. GINA scores were assessed from asthmatic subjects. Each participant provided written informed consent. The study was approved by the local ethics committee (5534/12). Among treated AR patients, 9 patients were additionally affected by asthma, 10 among intreated. Asthmatic patients are represented as a subgroup of the rhinitis patients, if not otherwise indicated. Patients were considered as asthmatic based on previous physician’s diagnosis and with a reported history of shortness of breath, cough, chest tightness during natural allergen exposure and/or earlier documented positive bronchodilation test. All patients were in good health (FEV1% >70%) with a history of clinically significant hay fever during the grass-pollen season since more than two years. Patients of the AIT-treated groups received at least one year of AIT treatment. Thereby, grass-pollen allergic patients with a history of moderate-severe and chronic persistent allergic rhinitis as defined by ARIA (Allergic Rhinitis and its Impact on Asthma) criteria since >2 years during the grass-pollen season, a positive skin prick test wheal >3mm in diameter and grass pollen specific IgE-level above 0.70kU/L underwent subcutaneous grass-pollen AIT.

In addition, total IgE was measured from sera of all individuals included in this study. Peripheral blood samples from all subjects included in this study were drawn at the same time points as sputum samples were analyzed by flow cytometry. For flow cytometric analysis, CPT tubes (BD biosciences) were centrifuged according to manufacturer’s instructions. Further, each sample was adjusted to 2.0×10^6^ cells and used for subsequent FACS staining. All procedures were performed in the Allergy Section, Department of Otolaryngology, TUM School of Medicine, Munich, Germany.

### Sputum collection, processing and characterization

Collection and processing of sputum as well as differential cell counts was performed as previously described ^48^. Briefly, human participants first inhaled salbutamol and consecutively nebulized hypertonic saline at increasing concentrations of 3%, 4%, and 5% NaCl every 7 min. During this procedure, participants cleaned their noses and rinsed their mouth to reduce squamous epithelium cells in the samples. Sputum was processed within one hour of collection. The selected sputum plugs, which contained as little saliva as possible were placed in a weighed Eppendorf tube and processed with 4x weight/volume of sputolysin working solution (Merck). Afterwards, 2x weight/volume of PBS was added. Samples were filtered through a 70 μm mesh and centrifuged for 10 min at 790 x g without break to remove the cells. Supernatants were stored at −80°C until further analysis. In addition, sputum cell slides were prepared for differential cell counts. Sputum samples were successfully collected from healthy controls (n=24; 92.3%) and allergic patients (n=34; 89.4%) once in and out of grass pollen season as previously described ^48^ and analyzed for secreted protein levels of TGF-β, IL-9, type-2 cytokines IL-4, IL-5, and IL-13, anti-inflammatory cytokine IL-10, type-1 cytokines IL-2, IFN-γγ and the pro-inflammatory cytokine IL-6. Cytokine levels were analyzed by LegendPlex TGF-β 1 and multiplex human inflammation panel assay (Biolegend). All nine parameters were detectable in every sputum sample derived from patients with AR and AA and compared to healthy control subjects.

### Lung function testing in humans

Baseline lung function was evaluated using a calibrated handheld pulmonary function testing device (Jaeger SpiroPro). The following parameters were recorded: vital capacity (VC), forced expiratory volume (FEV1), FEV1/VC, and maximum expiratory flow 25% (MEF 25%). Bronchodilator reversibility was tested after 400 μg of salbutamol.

### Total serum IgE measurement

Total serum IgE was assessed using diagnostic ImmunoCAP assays on a Phadia 100 device.

### Flow Cytometric Analysis of human and murine samples

Human sputum cell or PBMCs samples were labeled without stimulation for flow cytometry with specific antibodies using the Foxp3/Transcription Factor Staining Buffer Set (eBioscience) according to manufacturer’s instructions.

For flow cytometric analyses of the murine samples, staining for transcription factors was performed by using Foxp3 Staining Buffer Set (eBiosciences), while cytokine staining was performed by using the fixation/permeabilisation solution Kit (BD Bioscience Cytofix/Cytoperm™) according to the manufacturer’s protocols. For intracellular cytokine staining cells were stimulated for 4h with 50 ng/ml PMA (Applichem), 500 ng/ml Ionomycin (Thermo Fisher Scientific) and 1:1000 GolgiPlug (BD Bioscience). Flow cytometric analysis was performed using a BD LSRII Fortessa flow cytometer (BD Bioscience). Flow cytometry data were analyzed with FlowJo software (FlowJo). For gating strategy of human sputum cells (0.5×10^6^ cells) or PBMCs (2.0×10^6^ cells) samples see Fig. 5A and Fig. S2B; the murine gating strategy is shown in Fig.S1. Antibodies used for flow cytometry are listed in Table S2 (murine) and Table S6 (human).

### Real-time polymerase chain reaction

Total RNA was extracted either from mouse total lung tissue after homogenization using the RNeasy Mini Kit (Qiagen GmbH) or from enriched mouse lung CD4^+^ T cells (CD4 T cell isolation kit, Miltenyi Biotec, Auburn, CA, USA), using RNeasy Micro Kit (Qiagen) according to supplier’s instructions. RNA was reverse-transcribed directly (RevertAid H Minus First Strand cDNA Synthesis Kit, Thermo Scientific) and quantitative real-time PCR was performed using SYBR Green PCR Kit Master Mix (Qiagen) and the LightCycler^®^480 System (Roche) as previously described ^49^. The used primer sequences are listed in table S3. Each reaction was performed in duplicate in 284-well plates (Applied Biosystems) and turned into mean. The expression levels were normalized to GAPDH house-keeping gene and relative changes were represented as 2^−ΔΔCT^ (ΔΔC_T_=ΔC_T_−ΔC_Control_).

### Data acquisition and statistical analysis

All experimental procedures and analyses were conducted by blinded research staff. For the mouse experiments, differences between the groups in AHR were evaluated with two-way analysis of variance (ANOVA) with Bonferroni’s post-hoc test. Otherwise, differences between two data sets were evaluated using unpaired two-tailed Mann-Whitney test or t-test. Data were expressed as mean ± S.D., if not otherwise indicated. All statistically significant differences were depicted as *p<0.05, **p<0.01, ***p<0.001, ****p<0.0001. Data were analyzed using Prism software version 6 (GraphPad software Inc.). For the analysis of human samples, non-parametric statistical test were chosen, as the data points were not normally distributed. For figures 4 and 5, Kruskal-Wallis tests were performed initially to avoid multiple testing, and, only when medians across patient groups varied significantly, multiple single comparisons were performed using two-tailed Mann-Whitney U tests. Two-sided Spearman correlation was used to correlate IgE or RQLQ with immune cell frequencies in Figure 5.

## Acknowledgments

This study was supported by the German Center for Lung Research (DZL) to C.S.W. and U.M.Z., Helmholtz Inflammation&Immunology (I&I) to C.S.W., Grant of the German Research Foundation (DFG) No. 398577603 to C.S.W. and U.M.Z., and No. TR22 to T.B.

## Abbreviations

AA: allergic asthma
AAI: allergic airway inflammation
AHR: airway hyperreactivity
AIT: allergen-specific immunotherapy
AR: allergic rhinitis
BAL: Bronchoalveolar lavage
BALF: Bronchoalveolar lavage fluid
ICI: Inflammatory cell infiltrate
i.p.: intraperitoneal
Mch: metacholin
n.d.: not detectable
OVA: ovalbumin
PBMC: peripheral blood mononuclear cell
PBS: Phosphate-buffered saline
mRQLQ: Rhinoconjunctivitis Quality of Life mini Questionnaire
tIgE: total Immunoglobulin E
Tregs: regulatory T cells
Th: T helper cells

---

**Fig. S1.**
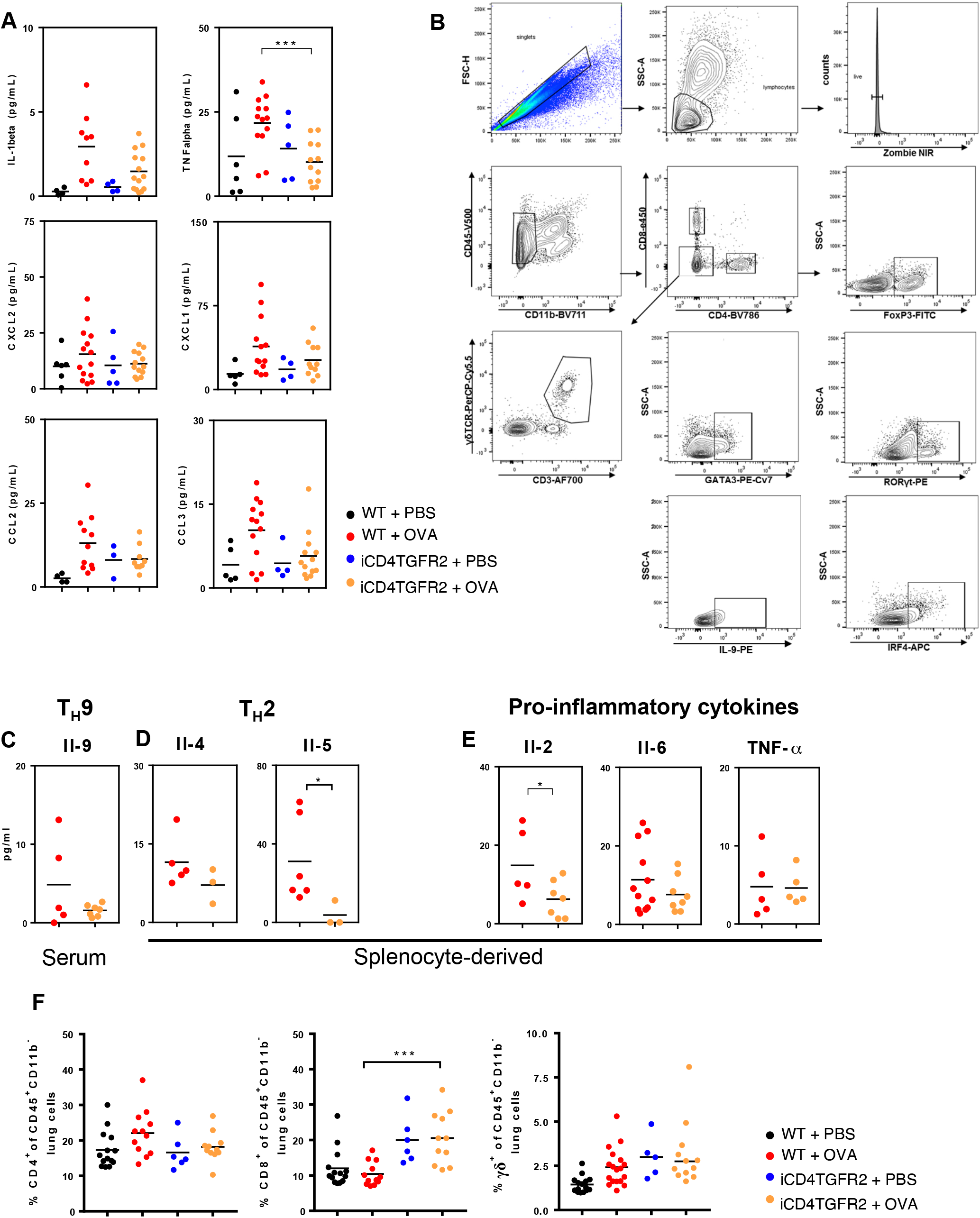

**Fig. S2.**
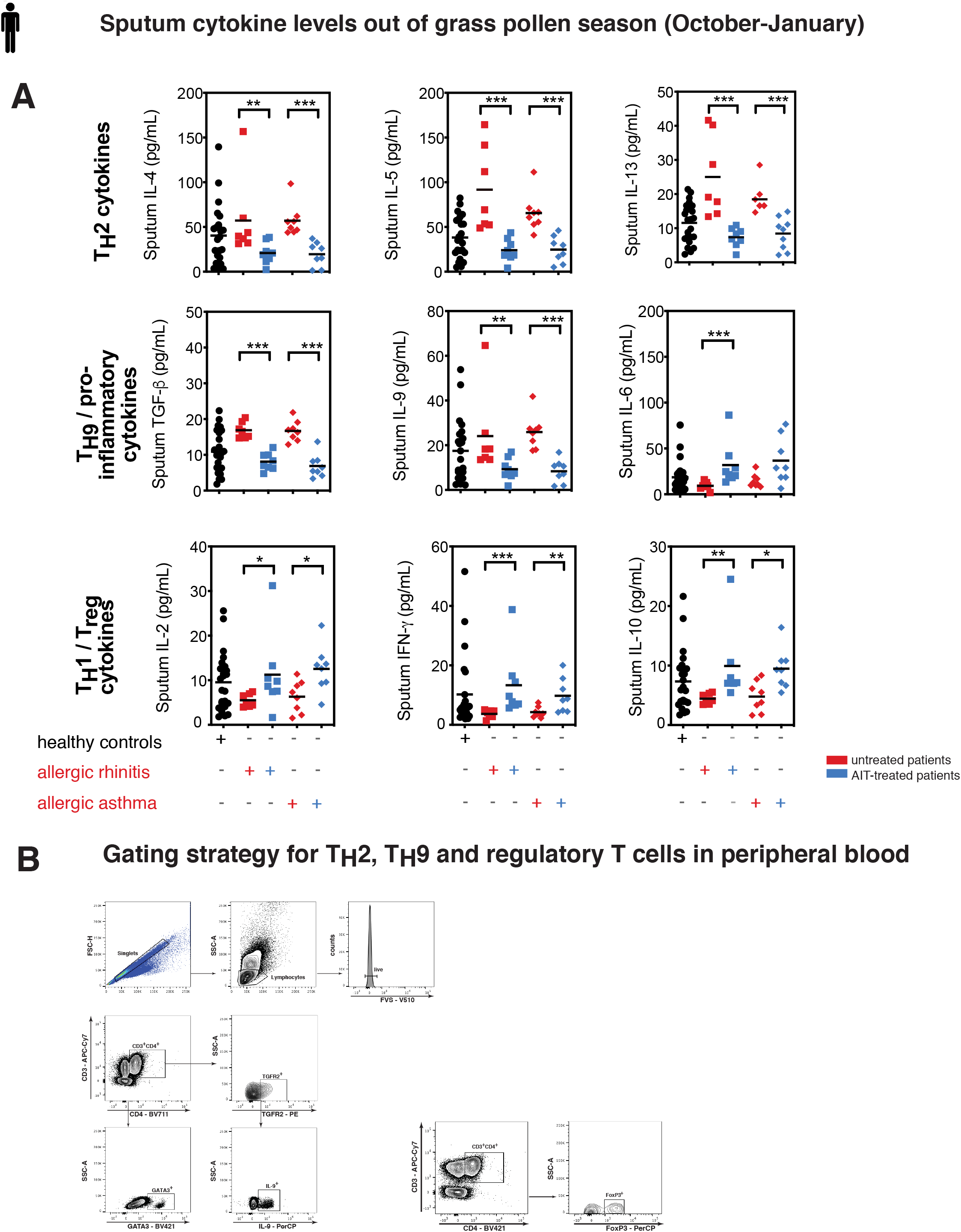

## References

1. Massague J. TGF-beta signal transduction. Annu Rev Biochem 1998; 67: 753–791.

2. Chang Z, Kishimoto Y, Hasan A, Welham NV. TGF-beta3 modulates the inflammatory environment and reduces scar formation following vocal fold mucosal injury in rats. Dis Model Mech 2014; 7(1): 83–91.

3. Putheti P. Polarizing Cytokines for Human Th9 Cell Differentiation. Methods Mol Biol 2017; 1585: 73–82.

4. Annunziato F, Cosmi L, Liotta F, Maggi E, Romagnani S. Human Th17 cells: are they different from murine Th17 cells? Eur J Immunol 2009; 39(3): 637–640.

5. Zhou L, Ivanov, II, Spolski R, Min R, Shenderov K, Egawa T et al. IL-6 programs T(H)-17 cell differentiation by promoting sequential engagement of the IL-21 and IL-23 pathways. Nat Immunol 2007; 8(9): 967–974.

6. Zissler UM, Chaker AM, Effner R, Ulrich M, Guerth F, Piontek G et al. Interleukin-4 and interferon-gamma orchestrate an epithelial polarization in the airways. Mucosal Immunol 2016; 9(4): 917–926.

7. Coker RK, Laurent GJ, Shahzeidi S, Lympany PA, du Bois RM, Jeffery PK et al. Transforming growth factors-beta 1, -beta 2, and -beta 3 stimulate fibroblast procollagen production in vitro but are differentially expressed during bleomycin-induced lung fibrosis. Am J Pathol 1997; 150(3): 981–991.

8. Coker RK, Laurent GJ, Shahzeidi S, Hernandez-Rodriguez NA, Pantelidis P, du Bois RM et al. Diverse cellular TGF-beta 1 and TGF-beta 3 gene expression in normal human and murine lung. Eur Respir J 1996; 9(12): 2501–2507.

9. Coker RK, Laurent GJ, Jeffery PK, du Bois RM, Black CM, McAnulty RJ. Localisation of transforming growth factor beta1 and beta3 mRNA transcripts in normal and fibrotic human lung. Thorax 2001; 56(7): 549–556.

10. Zhang M, Zhang Z, Pan HY, Wang DX, Deng ZT, Ye XL. TGF-beta1 induces human bronchial epithelial cell-to-mesenchymal transition in vitro. Lung 2009; 187(3): 187–194.

11. Ohno I, Nitta Y, Yamauchi K, Hoshi H, Honma M, Woolley K et al. Transforming growth factor beta 1 (TGF beta 1) gene expression by eosinophils in asthmatic airway inflammation. Am J Respir Cell Mol Biol 1996; 15(3): 404–409.

12. Vignola AM, Chanez P, Chiappara G, Merendino A, Zinnanti E, Bousquet J et al. Release of transforming growth factor-beta (TGF-beta) and fibronectin by alveolar macrophages in airway diseases. Clin Exp Immunol 1996; 106(1): 114–119.

13. Chu HW, Balzar S, Seedorf GJ, Westcott JY, Trudeau JB, Silkoff P et al. Transforming growth factor-beta2 induces bronchial epithelial mucin expression in asthma. Am J Pathol 2004; 165(4): 1097–1106.

14. Batra V, Musani AI, Hastie AT, Khurana S, Carpenter KA, Zangrilli JG et al. Bronchoalveolar lavage fluid concentrations of transforming growth factor (TGF)-beta1, TGF-beta2, interleukin (IL)-4 and IL-13 after segmental allergen challenge and their effects on alpha-smooth muscle actin and collagen III synthesis by primary human lung fibroblasts. Clin Exp Allergy 2004; 34(3): 437–444.

15. Chung Y, Lee SH, Kim DH, Kang CY. Complementary role of CD4+CD25+ regulatory T cells and TGF-beta in oral tolerance. J Leukoc Biol 2005; 77(6): 906–913.

16. Mantel PY, Schmidt-Weber CB. Transforming growth factor-beta: recent advances on its role in immune tolerance. Methods Mol Biol 2011; 677: 303–338.

17. Zissler UM, Jakwerth CA, Guerth FM, Pechtold L, Aguilar-Pimentel JA, Dietz K et al. Early IL-10 producing B-cells and coinciding Th/Tr17 shifts during three year grass-pollen AIT. EBioMedicine 2018; 36: 475–488.

18. Seumois G, Ramirez-Suastegui C, Schmiedel BJ, Liang S, Peters B, Sette A et al. Single-cell transcriptomic analysis of allergen-specific T cells in allergy and asthma. Sci Immunol 2020; 5(48).

19. Erpenbeck VJ, Hohlfeld JM, Discher M, Krentel H, Hagenberg A, Braun A et al. Increased expression of interleukin-9 messenger RNA after segmental allergen challenge in allergic asthmatics. Chest 2003; 123(3 Suppl): 370S.

20. Townsend JM, Fallon GP, Matthews JD, Smith P, Jolin EH, McKenzie NA. IL-9-deficient mice establish fundamental roles for IL-9 in pulmonary mastocytosis and goblet cell hyperplasia but not T cell development. Immunity 2000; 13(4): 573–583.

21. Gounni AS, Hamid Q, Rahman SM, Hoeck J, Yang J, Shan L. IL-9-mediated induction of eotaxin1/CCL11 in human airway smooth muscle cells. J Immunol 2004; 173(4): 2771–2779.

22. Dardalhon V, Awasthi A, Kwon H, Galileos G, Gao W, Sobel RA et al. IL-4 inhibits TGF-β beta-induced Foxp3+ T cells and, together with TGF-beta, generates IL-9+ IL-10+ Foxp3(−) effector T cells. Nat Immunol 2008; 9(12): 1347–1355.

23. Xue G, Jin G, Fang J, Lu Y. IL-4 together with IL-1beta induces antitumor Th9 cell differentiation in the absence of TGF-beta signaling. Nat Commun 2019; 10(1): 1376.

24. Tsuda M, Hamade H, Thomas LS, Salumbides BC, Potdar AA, Wong MH et al. A role for BATF3 in TH9 differentiation and T-cell-driven mucosal pathologies. Mucosal Immunol 2019; 12(3): 644–655.

25. Buttrick TS, Wang W, Yung C, Trieu KG, Patel K, Khoury SJ et al. Foxo1 Promotes Th9 Cell Differentiation and Airway Allergy. Sci Rep 2018; 8(1): 818.

26. Micosse C, von Meyenn L, Steck O, Kipfer E, Adam C, Simillion C et al. Human “TH9” cells are a subpopulation of PPAR-gamma(+) TH2 cells. Sci Immunol 2019; 4(31).

27. Mangan PR, Harrington LE, O’Quinn DB, Helms WS, Bullard DC, Elson CO et al. Transforming growth factor-beta induces development of the T(H)17 lineage. Nature 2006; 441(7090): 231–234.

28. Schwartz DM, Farley TK, Richoz N, Yao C, Shih HY, Petermann F et al. Retinoic Acid Receptor Alpha Represses a Th9 Transcriptional and Epigenomic Program to Reduce Allergic Pathology. Immunity 2019; 50(1): 106–120 e110.

29. Shull MM, Ormsby I, Kier AB, Pawlowski S, Diebold RJ, Yin M et al. Targeted disruption of the mouse transforming growth factor-beta 1 gene results in multifocal inflammatory disease. Nature 1992; 359(6397): 693–699.

30. Shimbara A, Christodoulopoulos P, Soussi-Gounni A, Olivenstein R, Nakamura Y, Levitt RC et al. IL-9 and its receptor in allergic and nonallergic lung disease: increased expression in asthma. J Allergy Clin Immunol 2000; 105(1 Pt 1): 108–115.

31. Cheng G, Arima M, Honda K, Hirata H, Eda F, Yoshida N et al. Anti-interleukin-9 antibody treatment inhibits airway inflammation and hyperreactivity in mouse asthma model. Am J Respir Crit Care Med 2002; 166(3): 409–416.

32. Sledzinska A, Hemmers S, Mair F, Gorka O, Ruland J, Fairbairn L et al. TGF-beta signalling is required for CD4(+) T cell homeostasis but dispensable for regulatory T cell function. PLoS Biol 2013; 11(10): e1001674.

33. Yee AA, Yin P, Siderovski DP, Mak TW, Litchfield DW, Arrowsmith CH. Cooperative interaction between the DNA-binding domains of PU.1 and IRF4. J Mol Biol 1998; 279(5): 1075–1083.

34. Chang HC, Sehra S, Goswami R, Yao W, Yu Q, Stritesky GL et al. The transcription factor PU.1 is required for the development of IL-9-producing T cells and allergic inflammation. Nat Immunol 2010; 11(6): 527–534.

35. Gomez-Rodriguez J, Meylan F, Handon R, Hayes ET, Anderson SM, Kirby MR et al. Itk is required for Th9 differentiation via TCR-mediated induction of IL-2 and IRF4. Nat Commun 2016; 7: 10857.

36. Mudter J, Yu J, Zufferey C, Brustle A, Wirtz S, Weigmann B et al. IRF4 regulates IL-17A promoter activity and controls RORgammat-dependent Th17 colitis in vivo. Inflamm Bowel Dis 2011; 17(6): 1343–1358.

37. Takami M, Love RB, Iwashima M. TGF-beta converts apoptotic stimuli into the signal for Th9 differentiation. J Immunol 2012; 188(9): 4369–4375.

38. Jones CP, Gregory LG, Causton B, Campbell GA, Lloyd CM. Activin A and TGF-beta promote T(H)9 cell-mediated pulmonary allergic pathology. J Allergy Clin Immunol 2012; 129(4): 1000–1010 e1003.

39. Tong R, Xu L, Liang L, Huang H, Wang R, Zhang Y. Analysis of the levels of Th9 cells and cytokines in the peripheral blood of mice with bronchial asthma. Exp Ther Med 2018; 15(3): 2480–2484.

40. Malik S, Sadhu S, Elesela S, Pandey RP, Chawla AS, Sharma D et al. Transcription factor Foxo1 is essential for IL-9 induction in T helper cells. Nat Commun 2017; 8(1): 815.

41. Wagner C, Uliczka K, Bossen J, Niu X, Fink C, Thiedmann M et al. Constitutive immune activity promotes JNK- and FoxO-dependent remodeling of Drosophila airways. Cell Rep 2021; 35(1): 108956.

42. Nouri-Aria KT, Pilette C, Jacobson MR, Watanabe H, Durham SR. IL-9 and c-Kit+ mast cells in allergic rhinitis during seasonal allergen exposure: effect of immunotherapy. J Allergy Clin Immunol 2005; 116(1): 73–79.

43. Kawayama T, Matsunaga K, Kaku Y, Yamaguchi K, Kinoshita T, O’Byrne PM et al. Decreased CTLA4(+) and Foxp3(+) CD25(high)CD4(+) cells in induced sputum from patients with mild atopic asthma. Allergol Int 2013; 62(2): 203–213.

44. Akdis M, Verhagen J, Taylor A, Karamloo F, Karagiannidis C, Crameri R et al. Immune responses in healthy and allergic individuals are characterized by a fine balance between allergen-specific T regulatory 1 and T helper 2 cells. J Exp Med 2004; 199(11): 1567–1575.

45. Alessandrini F, Schulz H, Takenaka S, Lentner B, Karg E, Behrendt H et al. Effects of ultrafine carbon particle inhalation on allergic inflammation of the lung. J Allergy Clin Immunol 2006; 117(4): 824–830.

46. Herz U, Braun A, Ruckert R, Renz H. Various immunological phenotypes are associated with increased airway responsiveness. Clin Exp Allergy 1998; 28(5): 625–634.

47. Wimmer M, Alessandrini F, Gilles S, Frank U, Oeder S, Hauser M et al. Pollen-derived adenosine is a necessary cofactor for ragweed allergy. Allergy 2015; 70(8): 944–954.

48. Zissler UM, Jakwerth CA, Guerth F, Lewitan L, Rothkirch S, Davidovic M et al. Allergen-specific immunotherapy induces the suppressive secretoglobin 1A1 in cells of the lower airways. Allergy 2021.

49. Marzaioli V, Aguilar-Pimentel JA, Weichenmeier I, Luxenhofer G, Wiemann M, Landsiedel R et al. Surface modifications of silica nanoparticles are crucial for their inert versus proinflammatory and immunomodulatory properties. Int J Nanomedicine 2014; 9: 2815–2832.

50. Zhang M, Chen H, Liu MS, Zhu KY, Hao Y, Zhu DL et al. Serum- and glucocorticoid-inducible kinase 1 promotes insulin resistance in adipocytes via degradation of insulin receptor substrate 1. Diabetes Metab Res Rev 2021; 37(4): e3451.

